# Inter-microscope comparability of dental microwear texture data obtained from different optical profilometers

**DOI:** 10.1101/2022.03.08.483539

**Authors:** Daniela E. Winkler, Mugino O. Kubo

## Abstract

Dental microwear texture analysis has become a well-established approach for dietary inference and reconstruction of mammals and other tetrapods, both extant and extinct. As the amount of available data grows immensely, researchers could benefit from combining data gathered by others to perform meta-analyses. However, different devices used to capture three-dimensional surface scans for DMTA are known to produce variation even when measuring the same surface. Here we compare DMTA data of 36 guinea pigs that received different diets in a controlled feeding experiment, measured on a confocal disc-scanning and a confocal laser-scanning microscope. We are testing different pre-analysis filtering protocols to mitigate differences. We find inter-microscopes and filter-related differences for the majority of analysed 41 DMTA parameters. Certain microscope-specific filter routines resulted in less differences than other pre-analysis protocols. We further identify DMTA parameters which were stable regardless of microscope or pre-analysis treatment. Overall, the results obtained on both microscopes show the same dietary differentiation between guinea pig feeding groups, which supports that DMTA is a suitable method to obtain repeatable, objective dietary inferences. We finally propose a roadmap to enhance data exchange and inter-lab comparability and collaboration in the future.

## Introduction

The analysis of microscopic wear features on teeth to infer dietary preferences of extant and extinct vertebrates, as well as use-wear analysis to infer the function of man-made tools, is of great interest to a broad community of biologists, palaeontologists, and archaeologists. Methods have undergone distinct modifications since microwear analysis was developed Baker et al. (1959) and gained popularity since Walker et al. (1972), with a recent trend to capture three-dimensional surface data in semi-automated, repeatable, and less observer-biased approaches commonly summarized under the moniker dental microwear texture analysis (DMTA). Within DMTA, different algorithms are applied to quantify surface texture patterns according to standardized scale-sensitive fractal analysis (SSFA) (e.g., Ungar et al. 2003, 2007; Scott et al. 2005, 2012) or International Organization for Standardization (ISO) surface roughness parameters (Schulz et al. 2010, 2013). The market standard for parameter computation is currently the software MountainsMap (DigitalSurf, France). More recently, MountainsMap-specific surface parameters such as mean depth and density of furrows, mean height, mean area, or motif analysis were also found indicative of dietary preferences and are frequently included into DMTA (e.g., Schulz 2013; Schulz-Kornas et al. 2019; Winkler et al. 2019a, b, 2020, 2021).

DMTA is relatively easily accessible through interferometry, confocal and laser-scanning microscopes of different price ranges, and several labs have established their own workflow, based on their specific devices. Thus, large amounts of data are being generated, and would likely enable meta-analyses if made available. However, the problem of availability is currently an important debate not only in the narrower DMTA community, but across disciplines. Most scientific journal encourage (or demand) all raw data to be made available through online repositories, but this leads to two problems:

1. Not all researchers agree to make raw data accessible at the moment of publication. This could be due to lax journal data availability policies, unwillingness to pay for storage services such as Dryad, or because they want exclusive access to their data for follow-up studies.
2. Data is being scattered across platforms, archives, and repositories (e.g., Zenodo, Dryad, github), which makes it hard to corroborate which data are available and which not.

The solution could be a joint initiative to store raw DMTA data, comparable to resources such as MorphoSource (www.morhposource.org) or the Paleobiology database (paleobiodb.org). However, if we achieve raw data availability, there is another problem to face:

How comparable are data obtained under different conditions? Data quality can potentially be affected on three levels. First, by whether scans have been obtained directly from original specimens (teeth, bones, artifacts) or from casts. Mihlbachler et al. (2019) found that scans obtained from casts gave significantly different DMTA results than scans obtained from original enamel surfaces, and that discrimination between dietary groups was thus diminished. Second, Goodall et al. (2015) highlighted not only differences between scans obtained from originals and casts, but also a second obstacle, that some moulding silicones are more accurate than others. It is therefore crucial to know how data were obtained (from original versus mould/cast), and to be aware of potential variation in fidelity of surface reproduction from different moulding compounds. Still, such methodological problems can be easily tackled by extending comparative studies of the widely available moulding materials and urging the community towards using the highest precision material available.

The third level of data quality differences, however, may potentially be the largest hindrance in making available data comparable. Data gathered from different confocal (blue and white light) profilers and laser-scanning microscopes has been found to differ for the same specimens. Even within the same product line (5 different Sensofar *PLμ* or 2 different Olympus LEXT), the same surface scan obtained on different microscopes produced different DMTA data (Arman et al. 2016, 2019). The authors found that differences could be reduced by using an automated pre-analysis filter-routine to mitigate measurement noise. Similarly, for two confocal laser-scanning microscopes of the same product line (Keyence VK), Kubo et al. (2017) found that application of different pre-analysis filter routines had a stronger effect on DMTA data than the microscope used. These are, up to now, the only studies comparing inter-microscope differences when generating DMTA data from biological surfaces. Besides Sensofar, Olympus, and Keyence, also confocal (laser and light) microscopes made by Zeiss (LSM 800) and NanoFocus (*μ*surf) are frequently used to obtain DMTA data. The different scanning methods and specifications (numerical aperture, spatial and lateral resolution etc.) will likely result in different results when scanning the same sample (Calandra et al. 2019a), a problem that has often been discussed by metrologists (see Arman et al. 2016 and references within). Moreover, manufacturers do not share how data is processed from capture to output, thus we cannot assume that data will be comparable when captured on these different devices. We can streamline post-scanning filtering protocols (and statistical approaches for analysis) and usage of moulding material, but we cannot easily understand and accommodate for the differences in data processing right after scanning, which is essentially a black box.

Consequently, if we want to achieve comparability, and make use of the massive amounts of data generated by the community in the future, we need to establish protocols to increase comparability and follow up on the approaches by Arman et al. (2016) and Kubo et al. (2017).

The aim of this study is to perform a cross-device, and cross-scanning method comparison by compiling an extensive dataset of the same individuals on both a confocal (blue) light microscope (*μ*surf Custom) and a confocal (violet) laser-scanning microscope (Keyence VK-9700). Both devices are equipped with a comparable 100x long-distance objective, but the scanning field differs. Spatial and vertical resolution and other specifications are similar between both devices (Table 1).

**Table 1.**
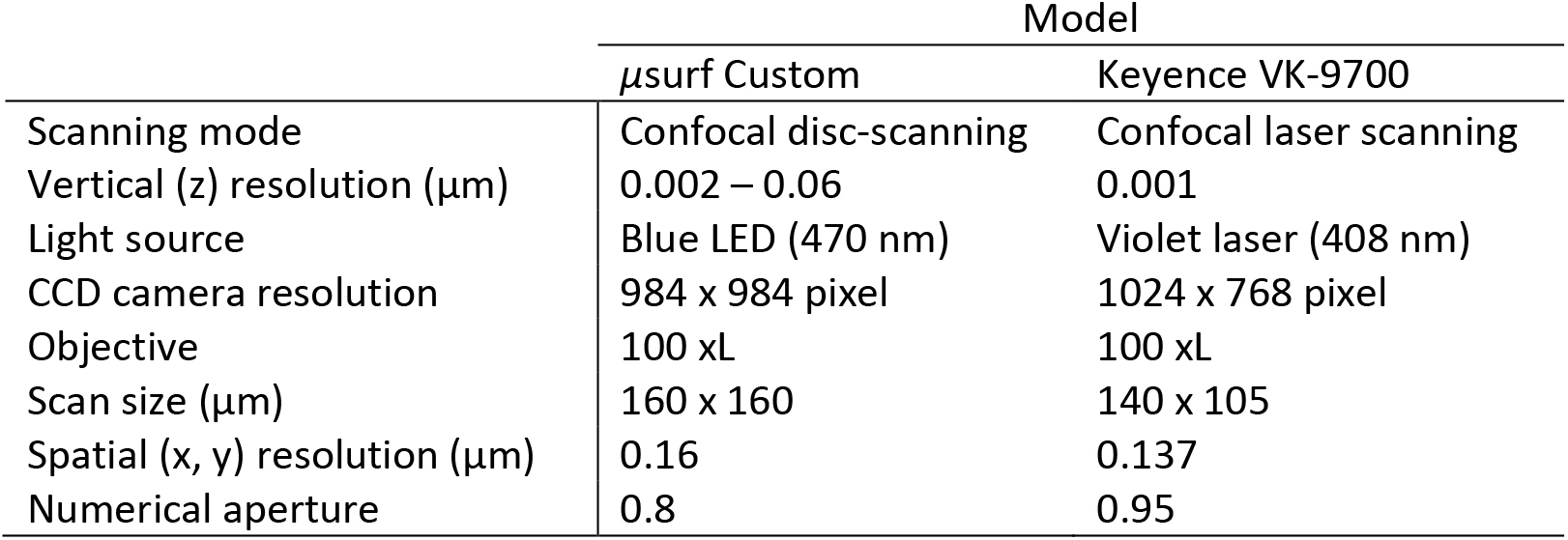
Technical specifications of the two confocal microscopes employed in this study.

We scanned the same dental moulds of 36 guinea pigs who received 6 homogenous diets in groups of 6 individuals during a 3-week feeding experiment (Winkler et al. 2019a, 2021), on both microscopes. Results are being compared in terms of relative differences between diet groups, but also in terms of absolute parameter values.

Under the given conditions, scanned areas are impossible to match perfectly when scanned on both devices, as the original scans were not taken with a repeated-measurement design in mind (see Boehm et al. 2019 and Calandra et al. 2019b for best practice how to prepare samples and the measuring system in order to match scans in a repeated-measurement design). Thus, we also test how well results of dietary differences can be reproduced when re-scanning the same individuals, and slightly varying the scanning positions.

## Material and Methods

Silicone moulds of the upper right dentition for 36 guinea pigs were made using high-resolution silicone (Provil novo Light C.D.2 fast set EN ISO 4823, type 3 light, Heraeus Kulzer GmbH, Dormagen, Germany). We decided to measure moulds, because the jugular bone would contact the microscope’s objective and obstruct the desired measuring position. The measurements took place almost 3 years apart, on the same silicone moulds. They were first scanned in spring 2018 on a *μ*surf Custom confocal disc-scanning microscope, in the Center of Natural History, University of Hamburg. Further details can be found in Winkler et al. (2019), the original raw surface scans are published in Winkler et al. (2021).

The second time, the moulds were scanned on a Keyence confocal laser-scanning microscope VK-9700, at The Graduate School of Frontier Sciences, University of Tokyo, in spring 2021. Technical specifications of both microscopes are given in Table 1.

### Description of scanning and filtering procedure

In both instances, the anteriormost enamel band of the upper right fourth premolar was scanned, and up to four non-overlapping scans taken. We were unable to match the exact same position of the previous scans. However, as the enamel band is narrow and short, it is very plausible that the re-scanned areas represent have a huge overlap with the previously scanned areas. Following Winkler et al. (2019), scans were then manually cropped in MountainsMap 8.2 to 60×60 μm, because enamel bands in guinea pigs are generally smaller in width than the default scanning areas (*μ*surf Custom: 160×160, Keyence VK-9700: 141×106um). The positioning of individual cut-outs will likely differ between scans obtained on *μ*surf Custom and Keyence VK-9700. As an exact matching cannot be ensured, the study also serves as a test of how well results can be reproduced when re-scanning the same individuals, and slightly varying the scanning positions.

Further data processing was conducted in MountainsMap 8.2, including re-analysis of the data obtained from the *μ*surf Custom at the University of Hamburg. Data were treated in two ways:

1. Applying a slightly modified version of the standard protocol established for the Keyence VK-9700 at the Kubo lab (Aiba et al. 2019, Kubo and Fujita 2021) which includes mirroring all surfaces in x and z (to compensate for the moulding procedure), levelling (least-square plane by subtraction), spatial filtering (robust Gaussian filter with a cut-off value of 0.8 μm), filling of non-measured points using the smoothing function of Mountains Map, noise-reduction by thresholding (upper and lower 0.5 %), removal of outliers (maximum slope of 85%) and form removal (polynomial of increasing power = 2). This protocol will hereafter be termed Filter A.
2. Following the standard protocol established for the *μ*surf Custom at the Kaiser Lab (Schulz et al. 2010, 2013) which includes mirroring all surfaces in x and z (to compensate for the moulding procedure), levelling (least-square plane by subtraction), spatial filtering (denoising median 5 x 5 filter size and Gaussian 3 x 3 filter size; default cut-offs are used), filling of non-measured points using the smoothing function of Mountains Map, noise-reduction by thresholding (upper and lower 0.5 %), removal of outliers (maximum slope of 85%) and form removal (polynomial of increasing power = 2). This protocol will hereafter be termed Filter B.

Both filtering protocols were applied for both datasets, resulting in four datasets:

Keyence*Filter A - *μ*surf*Filter A

Keyence*Filter A - *μ*surf*Filter B

Keyence*Filter B - *μ*surf*Filter A

Keyence*Filter B - *μ*surf*Filter B

We computed a total of 41 surface texture parameters that are frequently applied on biological surfaces (citations). There are some differences to the parameter set published in Winkler et al. (2019). We did not include the four ISO-12871 flatness parameters, as they are directly derived from the ISO-25178 height parameters *Sa*, *Sq*, *Sv* and *Sz* and thus redundant. Similarly, we excluded the ISO-25178 volume parameter *Vmp* because it is identical to *Vm* when using default cutoff settings. We additionally included the SSFA parameters *Asfc* (area-scale surface complexity) and *epLsar* (anisotropy) as these are among the most frequently applied measures of DMTA in other studies (Ungar et al. 2003, 2007; Scott et al. 2005, 2012; Merceron 2010; Schubert et al. 2010).

### Statistics

All statistical analyses were carried out in JMP Pro v.16. For each specimen, median values per parameter were calculated from up to 4 (at least 3) non-overlapping scans (compare Winkler et al. 2019a). Because of the repeated-measurement design, i.e. the same specimens were analysed four times through a combination of two microscopes and two filtering protocols, we performed a t-test for paired samples combined with a Wilcoxon signed-rank test to test for significant differences between the repeated measurements.

Additionally, we ranked data of each DMTA parameter within the same dietary group in order to standardize the difference between the dietary groups and applied a non-parametric Steel-Dwass test using the ranked dataset for all pairs.

## Results

Generally, absolute parameter values were shifted between both microscopes, but we found a good matching of dietary differences on both devices, and regardless of filtering protocols. The groups lucerne fresh, lucerne dry and grass fresh showed similar parameter values on both microscopes for most parameters (Fig. S1, Tab. S1). Grass dry fell between these three previous groups and the two remaining groups, bamboo fresh and bamboo dry.

Pairwise t-tests showed that all filtering routines performed very similar. The least significant differences between datasets were obtained for three filtering routines: either using Filter A on both datasets, or Keyence* Filter A and *μ*surf* Filter B, or Keyence*Filter B and *μ*surf*Filter A. Each resulted in 25 out of 41 significantly different parameters. The combination Keyence*Filter B and *μ*surf*Filter B showed 27 significantly different parameters. According to the Wilcoxon signed-rank test, differences between filtering routines were even smaller, with either 28 or 29 significantly different parameters reported (compare Tab. S2). The Steel-Dwass test on ranked data identified least significant differences for the filter combination Keyence* Filter A and *μ*surf* Filter B (20), followed by Keyence* Filter A and *μ*surf* Filter A (22) (Tab. S3).

The Keyence VK-9700 showed less outliers on for bamboo groups in height parameters. Generally, the Keyence VK-9700 produced slightly lower height and volume values on lucerne and fresh grass on than the μsurf Custom.

### Individual parameter groups

#### Area parameters

Data obtained using the *μ*surf Custom generally showed higher area values as data obtained on the Keyence VK-9700 (Tab. S1, Fig. S1). Least differences were found for *Sda* (standard dale area), which was the only not significantly different area parameter when using the filter routine Keyence*A – *μ*surf*B.

Both lucerne and the fresh grass group had the lowest area parameter values for both microscopes and filter combination. The dry grass group showed higher values than lucerne and fresh grass on *μ*surf Custom for all area parameter, and only for *Sda* on Keyence VK-9700. Both bamboo groups consistently showed the largest values for all area parameters. For data from *μ*surf Custom, the fresh bamboo group had higher values than the dry bamboo group, while for Keyence VK-9700 there was either no difference or dry bamboo had slightly higher values than fresh bamboo.

#### Complexity parameters

Complexity parameter values were generally higher when obtained on the Keyence VK-9700. While the shift was similar for all diet groups and the parameters *Sdr* and *Asfc*, the parameter *nMotif* (number of motifs) showed a larger offset for the dry grass group. Significant differences between microscopes were not detected for *Sdr* and *Asfc* when using the filter routine Keyence*A – *μ*surf*B.

For *Sdr* and Asfc, both lucerne groups and the fresh grass group had similarly low values on both microscopes. Dry lucerne showed slightly higher values when measured on the Keyence VK-9700. Dry grass and both bamboo groups showed higher values than the previous three groups. All three had comparable values when measured on *μ*surf custom (with fresh bamboo having highest variability), while on the Keyence VK-97000 dry grass and fresh bamboo had higher *Sdr* and *Asfc* values than dry bamboo. For *nMotif*, the dry grass group displayed strong differences between both microscopes. While on *μ*surf Custom it showed intermediate values between the lucerne groups (high) and the bamboo groups (low), it had higher values than all other diet groups when measured on VK-97000 (Figs. 1B, S1).

**Figure 1.**
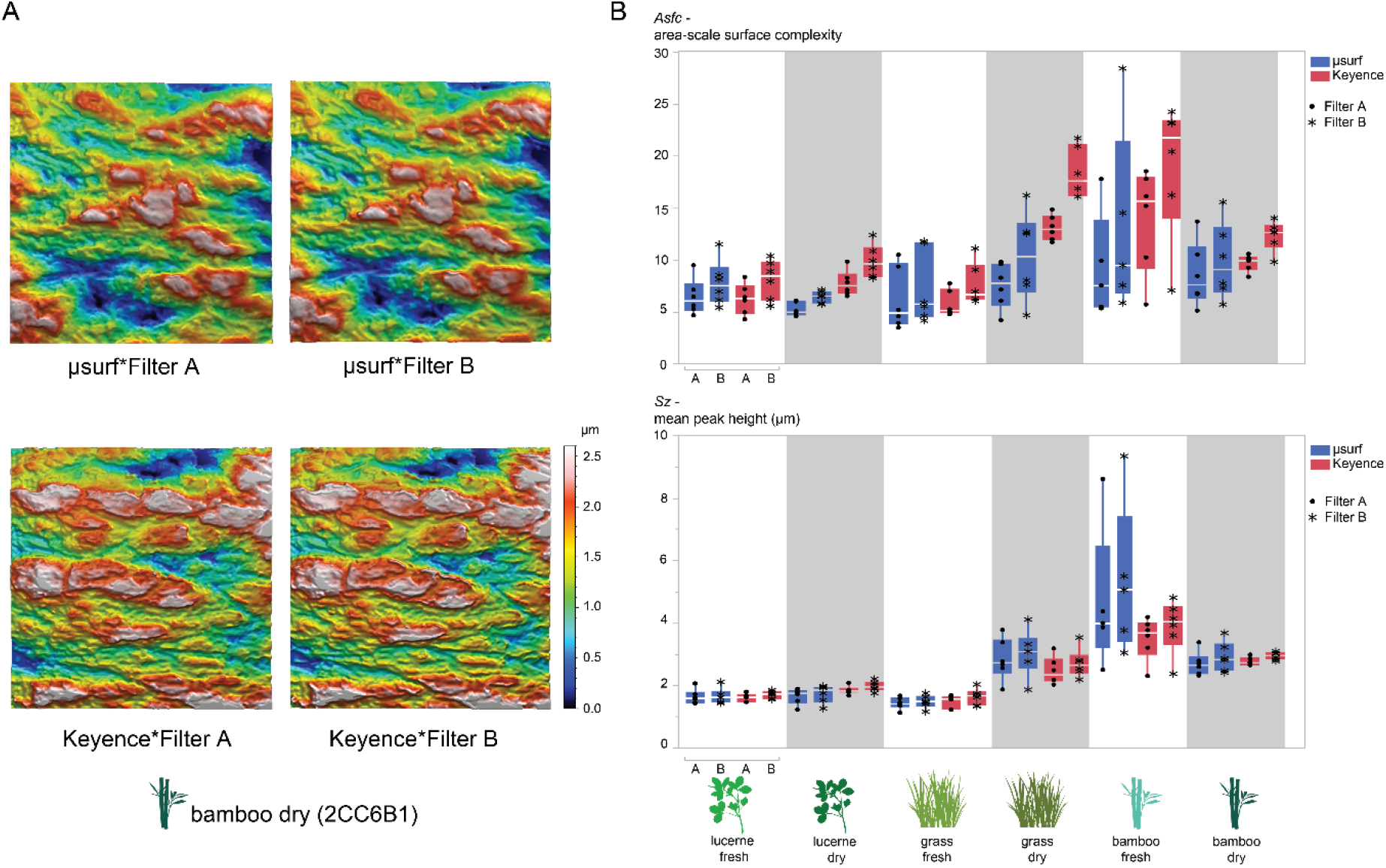
Comparison of data derived from either *μsurf Custom* or Keyence VK-9700. A) Exemplary 3D photosimulations of surfaces from one individual of the bamboo dry group, scanned on both microscopes and processed with Filter A and Filter B. Note that the scanning area does not match between microscopes. Scans are to the same scale (μm). B) Exemplary DMTA parameter (upper: *Asfc*, lower: *Sz*) calculated for all individuals, with data derived from *μ*surf Custom (blue) and Keyence VK-9700 (red) and different filter

#### Density parameters

While for *Sal* (autocorrelation length) the *μ*surf Custom produced higher values, for *Spd* (density of peaks) and *medf* (density of furrows), the Keyence VK-9700 recorded significantly higher parameter values. Significant differences could not be eliminated by any of the applied filtering routine combinations.

The general pattern of diet groups was maintained on both microscopes for the parameters *Sal* and *medf*. For the parameter *Spd* (Density of furrows), the dry grass group showed low parameter values like both bamboo groups when measured on the *μ*surf Custom, and high parameter values similar to both lucerne and the fresh grass group when measured on the Keyence VK-9700 (Fig. S1).

#### Direction parameters

Measures of absolute texture direction (*Std*, *Tr1R*, *Tr2R*, Tr3R) were highly variable on both microscopes and cannot be compared. For measures of isotropy (*Str*, *IsT*), the *μ*surf Custom showed higher values, and consequently for anisotropy (*epLsar*), lower values as compared to the Keyence VK-9700.

For the anisotropy and isotropy parameters (*Str*, *epLsar*, *IsT*), both microscopes showed the same general pattern of diet groups. The lucerne and grass groups showed higher isotropy (larger *Str*, *IsT*), while the bamboo groups showed larger anisotropy (larger *epLsar*) (Fig. S1).

#### Height parameters

Overall, the match between both microscopes was good for height parameters. Data from *μ*surf Custom shows higher variability for the Bamboo dry group than data captured on the Keyence VK-9700. Nine out of 14 height parameters were not significantly different for the filter combination Keyence*A – *μ*surf*A. For DMTA parameters with significant differences between the microscopes, data obtained on the *μ*surf Custom had higher values than data obtained on the Keyence VK-9700 (compare *S5v*, *Sa*, *Sq*, *Sv*, *Sxp*) (Figs. 1B, S1).

The general pattern of parameter values for the diet groups was the same on both microscopes and for all filter routines. Both lucerne groups and grass fresh had the lowest height parameter values, followed by dry grass with intermediate values, while both bamboo groups had the largest height parameter values (Fig. S1).

#### Peak sharpness

The only peak sharpness parameter *Spc* was significantly higher when measured on Keyence VK-9700. The divergence could not be corrected by any of the applied filter routines. The general pattern, however, was the same on both microscopes, with lucerne fresh/dry and fresh grass showing lower parameter values than dry grass and bamboo fresh/dry (Fig. S1).

#### Plateau size parameters

Values for both plateau size parameters were comparable, however, most filter routines resulted in significant differences between microscopes. Data obtained on the *μ*surf Custom were larger for *Smr* and *Smc* (except for the dry bamboo group).

The general parameter values pattern of all diet groups was the same for both microscopes and all filter routines (Fig. S1).

#### Slope

The one slope parameter *Sdq* was larger when measured on the Keyence VK-9700. By applying the filter routine Keyence*A – *μ*surf*B, the differences were no longer significant.

Both microscopes showed similar patterns for all diet groups (Fig. S1).

#### Volume

Volume parameters showed higher values for data from the *μ*surf Custom, with some exceptions for the dry bamboo group. Filtering reduced differences, with Keyence*A – *μ*surf*A being the best routine.

The general pattern of low volume parameter values for both lucerne groups and the fresh grass group, intermediate values for the dry grass group, and highest values for both bamboo groups was found on both microscopes and using all filter routines (Fig. S1).

#### Exemplary correction

By performing linear regression, it would be possible to obtain correction factors for each individual DMTA parameter, and thus facilitate comparison between microscopes. Exemplarily, the result of such a correction can be seen in Figure 2 for the parameter *medf* (mean density of furrows). Here, data obtained on Keyence VK-9700 (Filter A) has been corrected according to data obtained on *μ*surf Custom (Filter B) by the equation:

**Figure 2.**
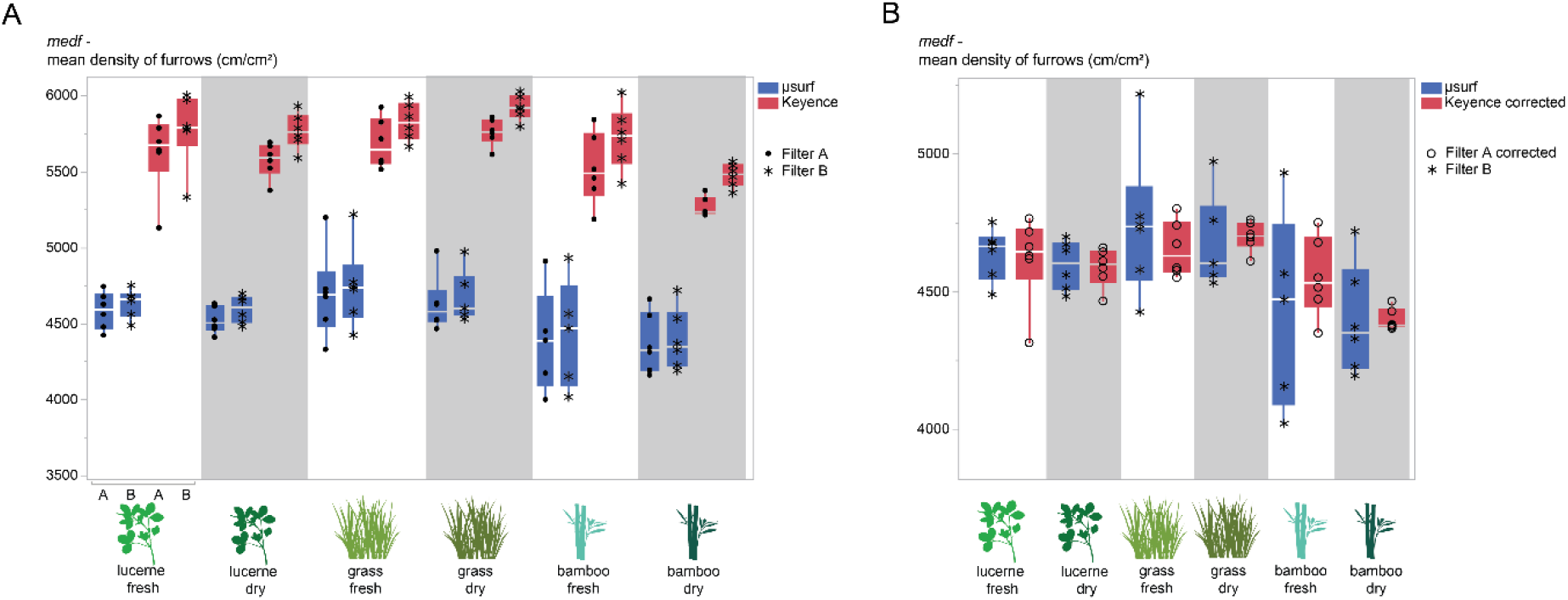
Comparison of results for the DMTA parameter *medf*. Scans were captured on either *μ*surf Custom (blue) or Keyence VK-9700 (red) and processed with different filter routines (Filter A: circle, Filter B: asterisk). A) All filter and microscope combinations. B) Data from Keyence*Filter A corrected according to linear regression equation and compared to *μ*surf*Filter B. Note that the data is well comparable after the correction.

*medf* corrected = 1151.6965 + *medf* from Keyence*0.6161424.

## Discussion

### Inter-microscope differences

Not all parameters are suitable for direct comparison of absolute parameter values when data is obtained on different devices. Our results clearly show that peak and furrow density-related parameters are significantly different and should not be used for immediate comparison. The source of this strong variation is likely due to the different scanning techniques and peak-detection algorithms of the two microscopes. Just by visually comparing, it is evident that scans from *μ*surf Custom appear slightly blurred, while scans from Keyence VK-9700 show sharper peaks (Figs. 1, S2). Through introduction of a correction factor, however, these microscope specific differences can be mitigated, and results become well comparable (Fig. 2).

Several DMTA parameters were similar in absolute values and could be compared with higher confidence (Tab. 2). Microscope-specific filtering protocols can thus help to minimize inter-microscope differences and account for device-specific characteristics. With the best filtering protocol, the congruence between data was 16 out of 41 (t-test) or 21 out of 41 (Steel-Dwass test) parameters, while the worst performing filtering protocol resulted in 14 or 15 out of 41 comparable parameters, respectively.

**Table 2.** Comparison of stable, i.e., not significantly different parameters, for the best performing filter combinations (based on t-test). Parameters that showed no significant differences regardless of filter protocol are given in bold. Directional parameters (*Std*, *Tr2R*, *Tr3R*) are interpreted as not diet-informative, but mostly related to chewing mechanics. Therefore, these are greyed out.

| Filter combination |  |  |
| --- | --- | --- |
| Keyence*A –<br>μsurf*A | Keyence*A –<br>μsurf*B | Keyence*B –<br>μsurf*A |
|  | Sda |  |
|  | Sdr |  |
|  | Asfc |  |
| Std | Std | Std |
| Tr2R | Tr2R | Tr2R |
| Tr3R | Tr3R | Tr3R |
| **S10z** | **S10z** | **S10z** |
| S5p | S5p |  |
| **Sku** | **Sku** | **Sku** |
| **Sp** | **Sp** | **Sp** |
|  |  | Sa |
|  |  | Sq |
| Ssk | Ssk |  |
| Sz |  | Sv |
|  |  | Sz |
| **meh** | **meh** | **meh** |
| madf |  | madf |
| **metf** | **metf** | **metf** |
|  |  | Smc |
| Smr | Smr |  |
|  | Sdq |  |
| Vm | Vm |  |
| Vv |  | Vv |
| Vvc |  | Vvc |

These results show that there is not one filter combination that yields unequivocally better results than the other. When considering results from all statistical tests, and for which parameters better comparability was achieved, we would favour the combination Keyence* Filter A and *μ*surf* Filter B. This conclusion is mainly based on the fact that frequently used parameters such as *Asfc*, *Sdr*, *Sda* and several height and volume parameters were comparable under this protocol. No other routine resulted in comparable results for the SSFA parameter *Asfc*, but as it is of great importance and generally applied, congruence should have high priority. Additionally, the applied filters were those originally developed for the specific microscopic devices (Schulz et al. 2010, 2013; Aiba et al. 2019; Kubo and Fujita 2021), which is additionally beneficial when considering data comparability of already published datasets.

### Comparability of results

The dietary differences between experimentally fed guinea pigs originally described by Winkler et al. (2019a) were confirmed for the newly obtained data from the Keyence confocal laser-scanning microscope, and by all filtering protocols. Both lucerne diets and the fresh grass diet were very similar in complexity, height, volume, area, and slope parameters. Especially low parameter values for height, volume, and the complexity parameters *Sdr* and *Asfc* are interpreted as related to low abrasive feeds (Kubo et al. 2017, Kubo and Fujita 2021; Schulz et al 2010, 2013; Winkler et al. 2019a, b, 2020). On the contrary, higher parameter values would indicate a more abrasive diet. Even though slightly different enamel areas were scanned, as matching of the original areas was not possible, the re-analysis gave the same result of increasing abrasiveness of experimental feeds in the order: Lucerne fresh, lucerne dry, grass fresh < grass dry < bamboo fresh, bamboo dry (Figs. 1B, S1). This shows that the previously derived interpretation of Winkler et al. (2019a) holds true, there is a significant effect of both silica content and hydration state of the plant tissue on observed microwear texture pattern, with dry and more siliceous diets resulting in more abrasion. Therefore, our study also provides a test for repeatability and reliability of DMTA results and strengthens the robusticity of the method.

### Outlook and suggested best practice

By finding the most stable (i.e., most comparable) parameters between the two microscopes, this study might help in the identification of 3D microwear texture parameters to focus on, as the huge availability has caused confusion and made it difficult to agree on a set of relevant parameters. By choosing the most stable ones, this controversy can be advanced. At least for comparative studies, where data is gathered in different labs, and on different machines, we suggest concentrating on the parameters listed in Table 2. The height parameters *S10z*, *Sku*, *Sp*, *meh* and *metf* did not show significant differences between microscopes, regardless of filter routines. Thus, we would consider these parameters as most stable and most suitable for comparative data analysis without adapted filtering routine, or introduction of a correction factor. For the parameters *nMotif* (number of motifs) and *Spd* (density of furrows), the dry grass group fell into a different position among the overall pattern when measured on the two different microscopes. Therefore, these parameters should be excluded from comparative analyses, and maybe also avoided generally, as the results were not reproductible. For parameters which showed a general shift, but could not be adjusted through filtering, a correction factor should be introduced as exemplarily shown here for the parameter *medf* (Fig. 2) if one desires to include them into a comparative study. Nevertheless, it must be noted that even after correction, results obtained from different microscopes need to be discussed as such, and the possibility of persisting inter-microscope differences needs to be discussed.

Such comparative studies should be conducted between more DMTA labs to understand specific characteristics of each microscope used, and to find best-practice filtering protocols that facilitate inter-microscope (and thus inter-lab) data comparability. This will lead to a massive increase of comparative data available for future studies and avoid unnecessary data-recollection. We highly encourage striving for a shared data repository to which DMTA research labs worldwide can contribute. As an initiative to promote such comparability, we propose to compile a standard set of moulds from different typical specimens (e.g., ungulate, reptile, carnivore) and a few standardized flat surfaces (e.g., polished enamel) that are mounted on a microtiter plate with incision marks that can be aligned within a microscope-specific coordinate system. Such specimens shall be exchanged between DMTA labs, with each research group re-scanning the same areas, and processing the data according to their own preferred pre-analysis protocol, and the published protocols of other researchers. Subsequently this data can be used to obtain accepted “correction equations” for each device, so that data can be shared and used between labs.

## Conclusion

Repeatability and less observer-biased interpretation of results are two key advantages often cited when comparing DMTA to classical microwear analysis. Our study supports these claims, as data gathered in two different laboratories, on two different microscopes, and 3 years apart resulted in the same dietary discrimination between experimentally fed guinea pigs.

This study also highlights that inter-microscope comparison can only be done without correction for a few DMTA parameter. The majority of often applied parameters needs to be corrected through a microscope-specific filter-routine, or a correction factor. Such correction factors could be obtained through a joint community effort which includes scanning of the same surfaces in multiple labs, which we propose here to our colleagues. Through our collaboration, we might achieve data comparability, and advance research in our field.

## Data availability

All original, unfiltered surface texture scans used in this study are available online. Data from Winkler et al. (2019a, 2021) has been published under <u>doi:10.25592/uhhfdm.9163</u>. The comparative scans obtained for this study are deposited under <u>doi.org/10.5061/dryad.7wm37pvw3</u>.

## Acknowledgements

This study was supported by grants-in-aid from the Japan Society for the Promotion of Science (JSPS), 20F20325 to DEW and no.16K18615 to MOK.

