## Supplementary material for "Inter-microscope comparability of dental microwear texture data obtained from different optical profilometers": Suppl. Figures

### Area

#### *Sda*

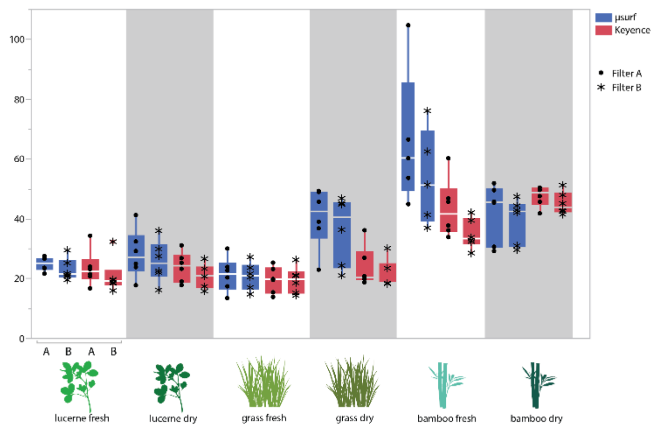


#### *Sha*

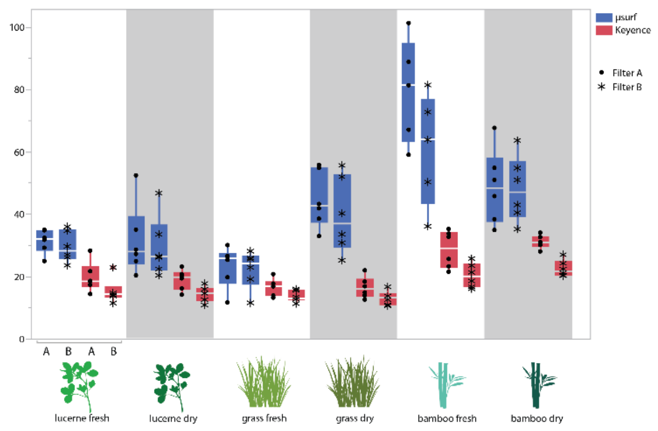


#### *mea*

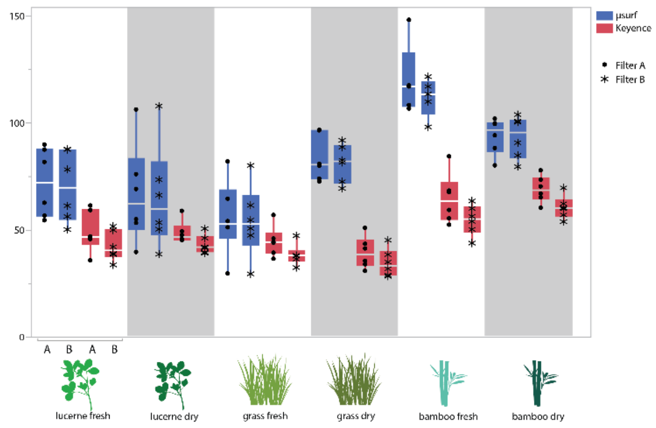

#### Complexity

*Sdr*

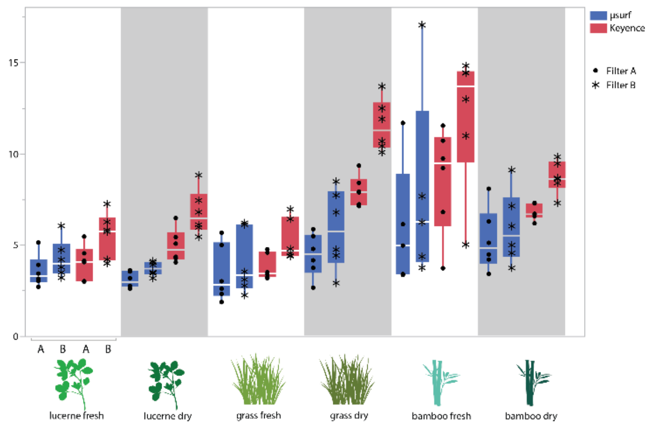


*Asfc*

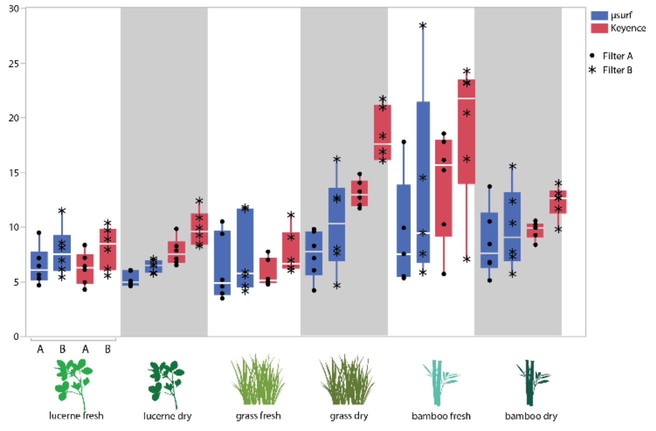


*nMotif*

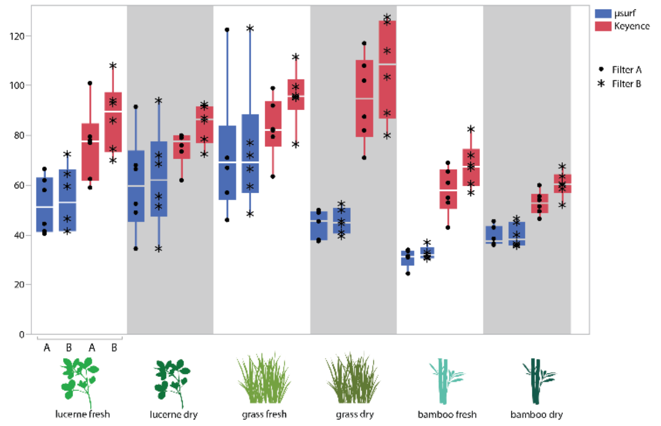

### Density

#### *Sal*

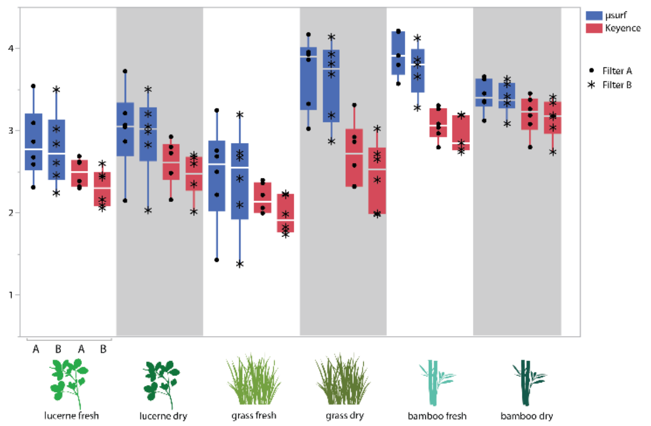


#### *Spd*

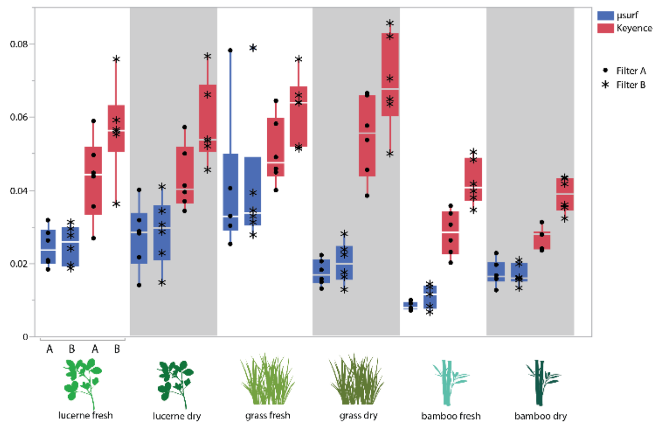


#### *medf*

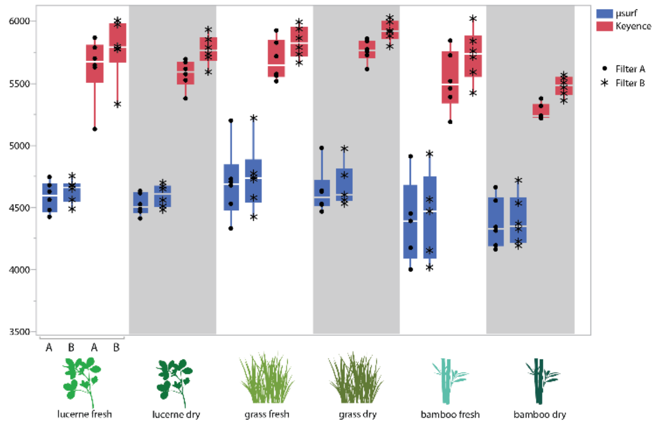


### Direction

#### *Std*

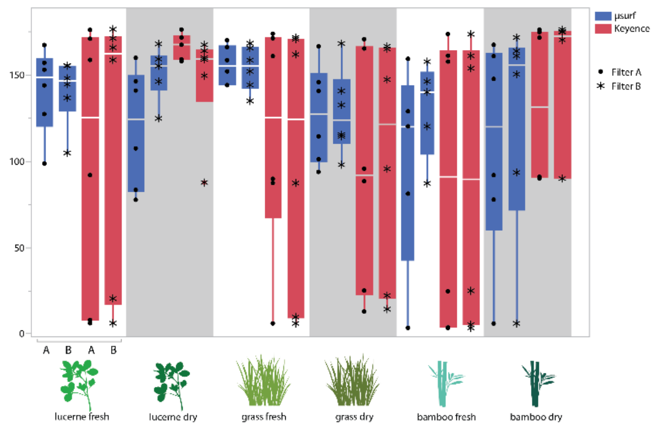


#### *Str*

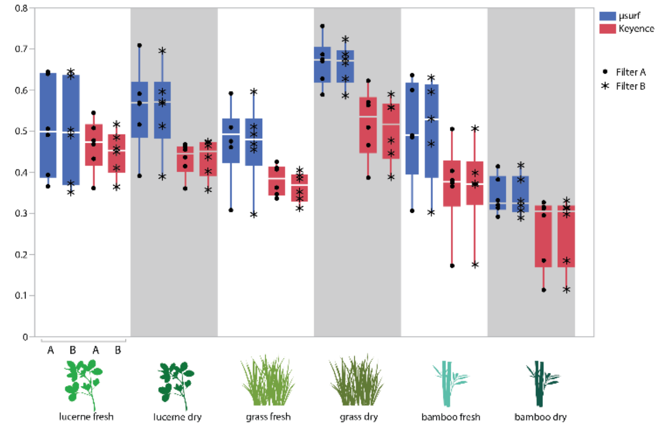

### Direction (cont.)

## *Tr1R*

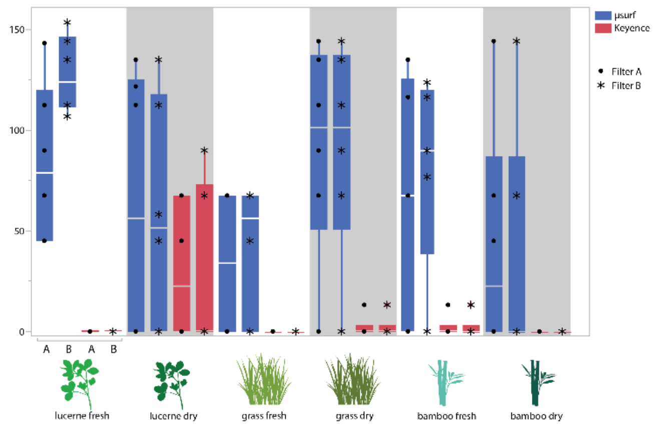


## *Tr2R*

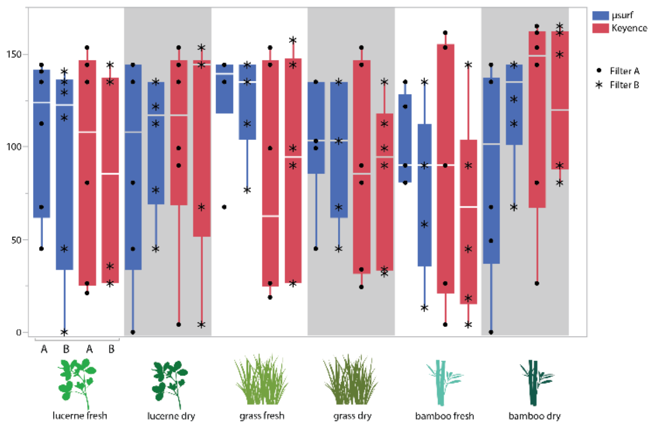


## *Tr3R*

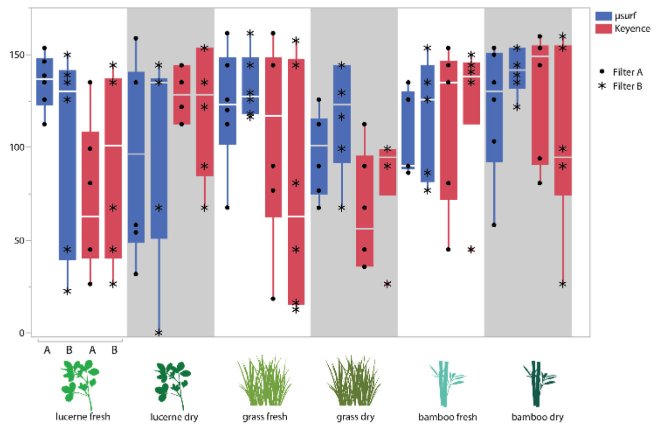


#### *epLsar*

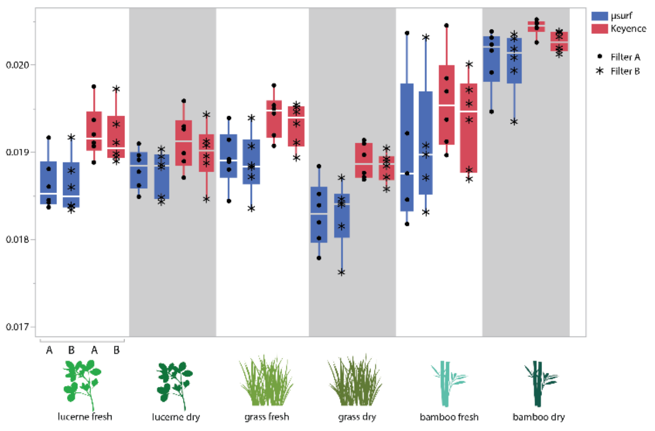


#### *IsT*

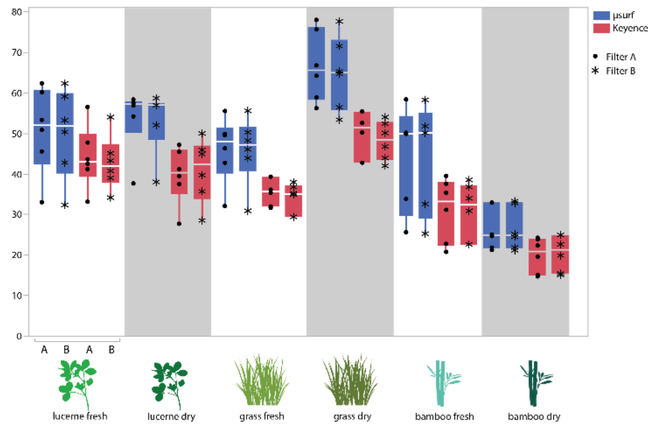

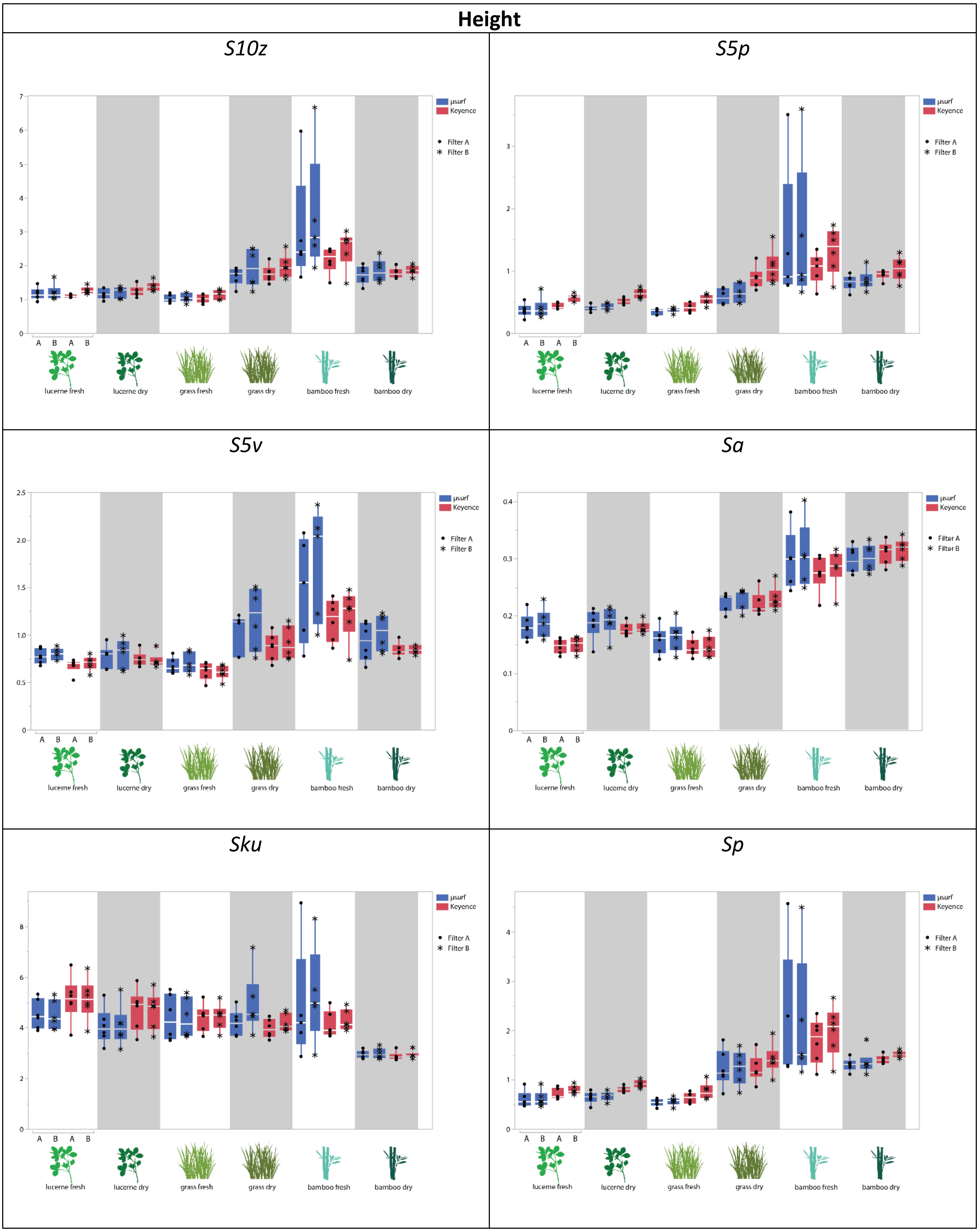
Height
S10z
S5p
S5v
Sa
Sku
Sp
µsurf
Keyence
Filter A
Filter B
A B A B
lucerne fresh
lucerne dry
grass fresh
grass dry
bamboo fresh
bamboo dry

### Height (cont.)

*Sq*

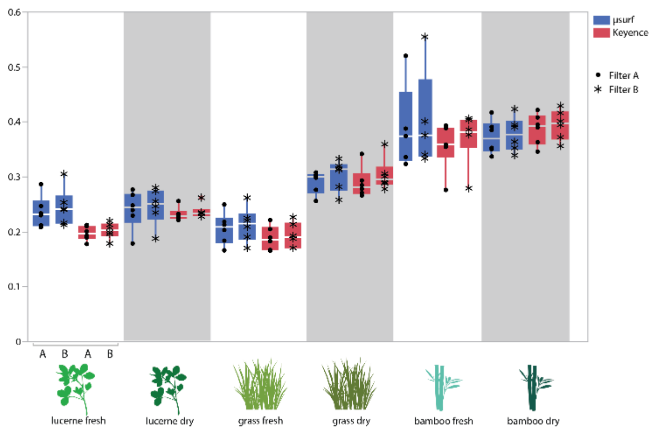


*Ssk*

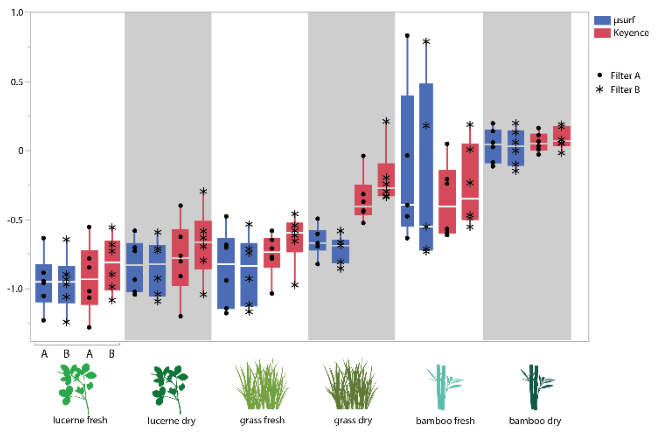


*Sv*

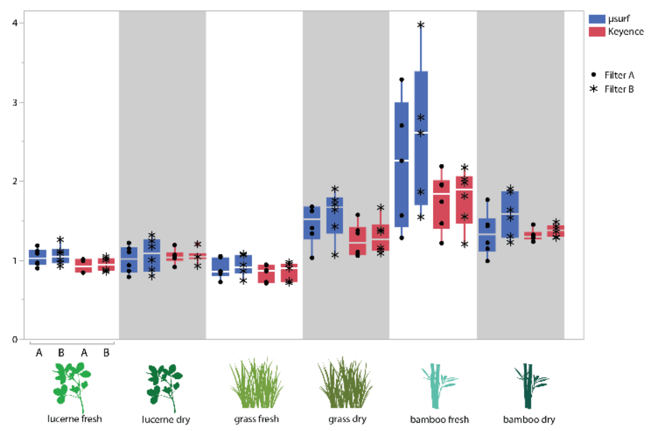


*Sxp*

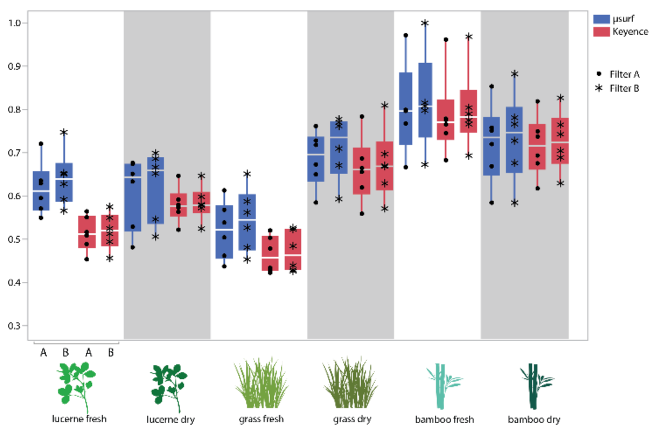


*Sz*

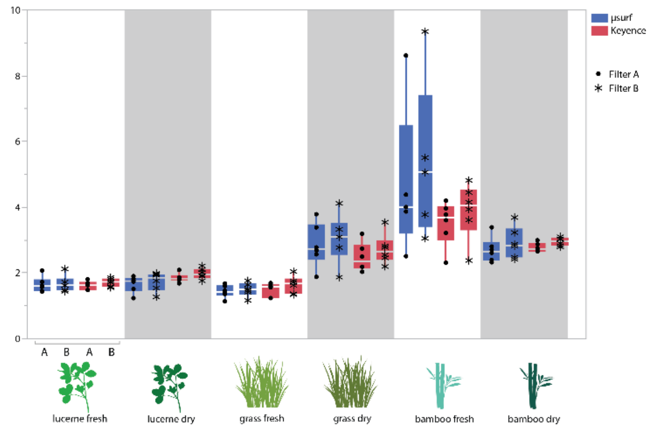


*meh*

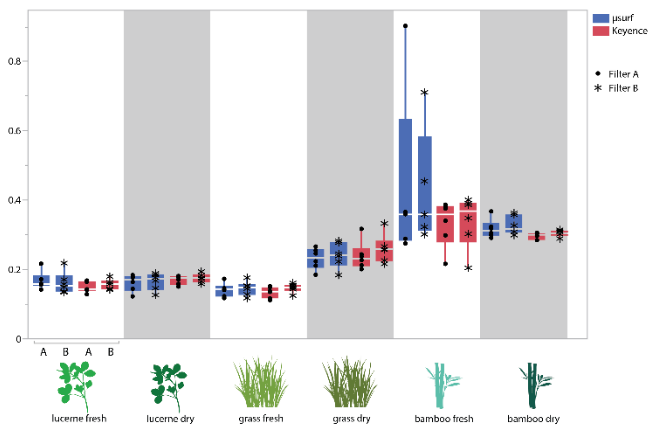

#### Height (cont.)

##### *madf*

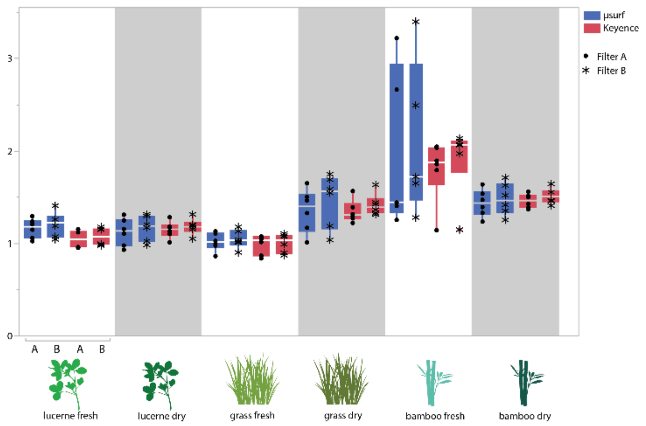


##### *metf*

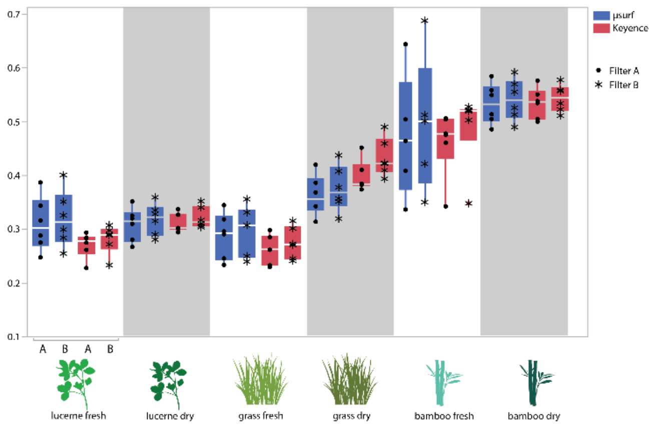


#### Peak sharpness

##### *Spc*

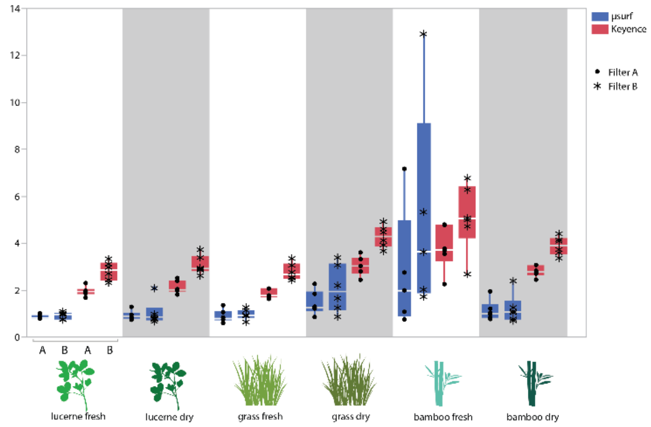


#### Plateau size

##### *Smc*

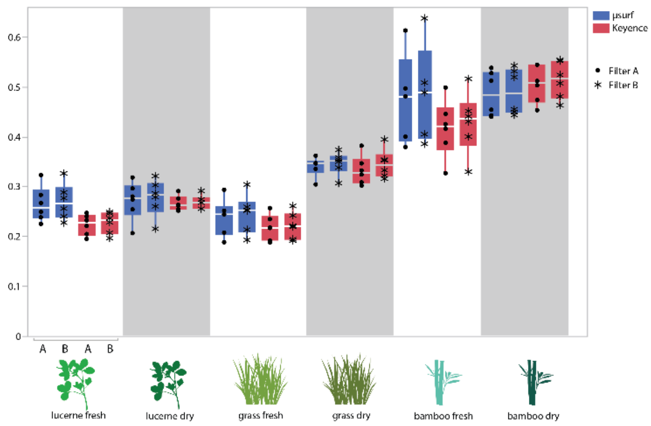


##### *Smr*

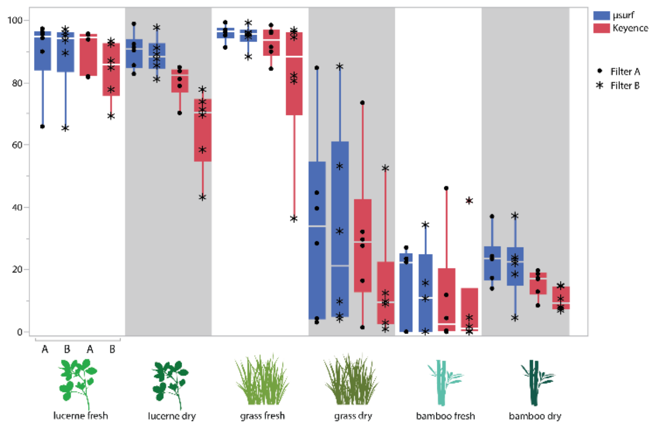

#### Slope

##### *Sdq*

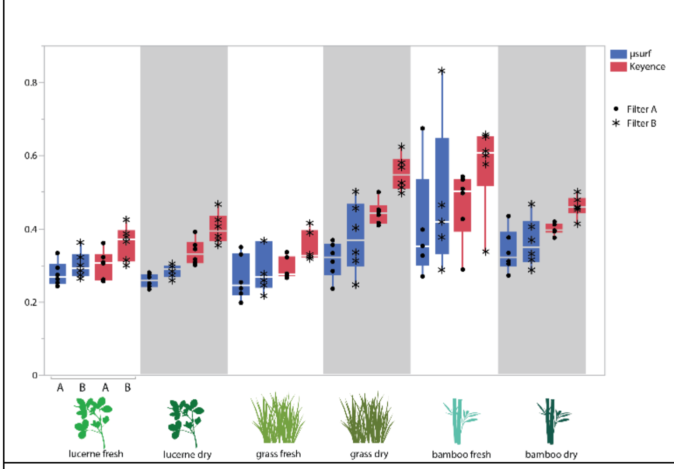


#### Volume

##### *Sdv*

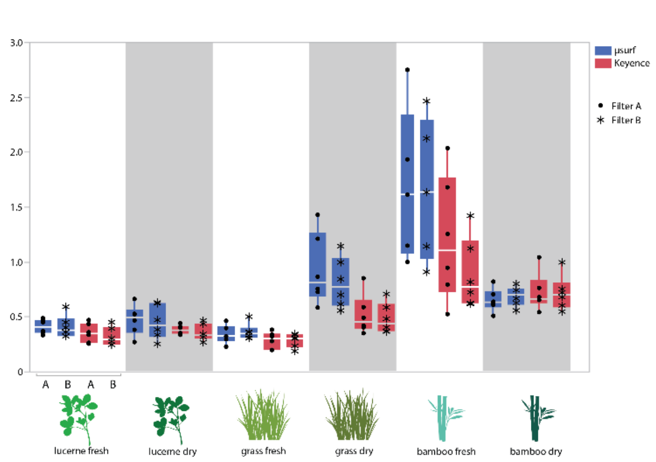


##### *Shv*

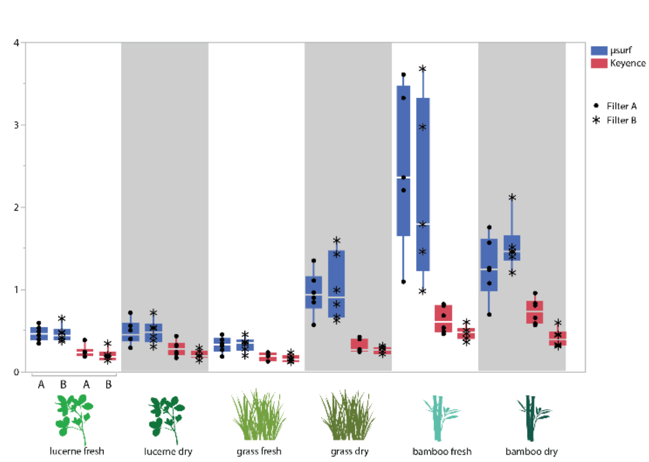


### *Vm*

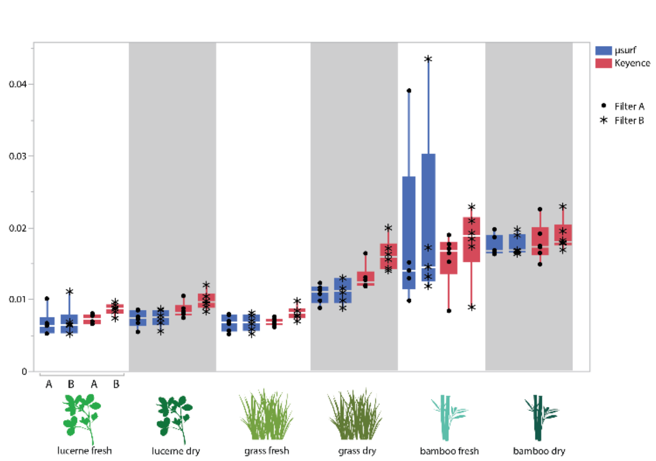


##### *Vmc*

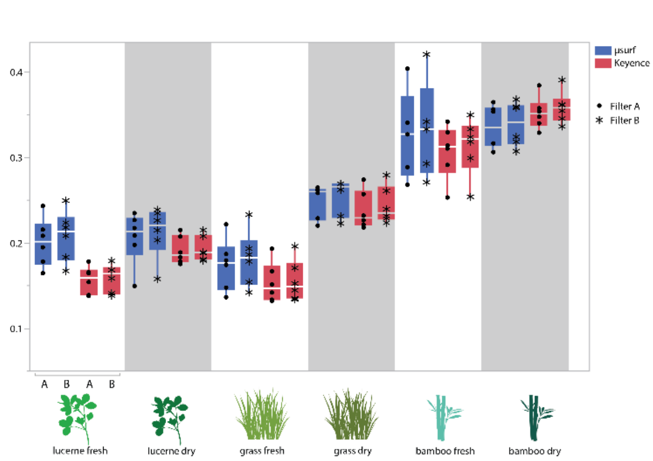

#### Volume (cont.)

*Vvc*

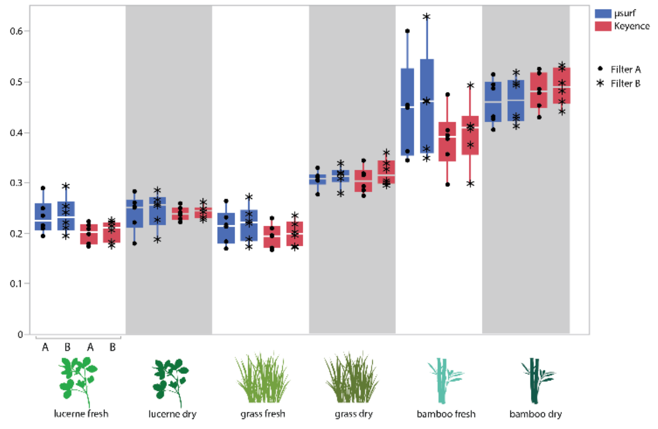


*Vv*

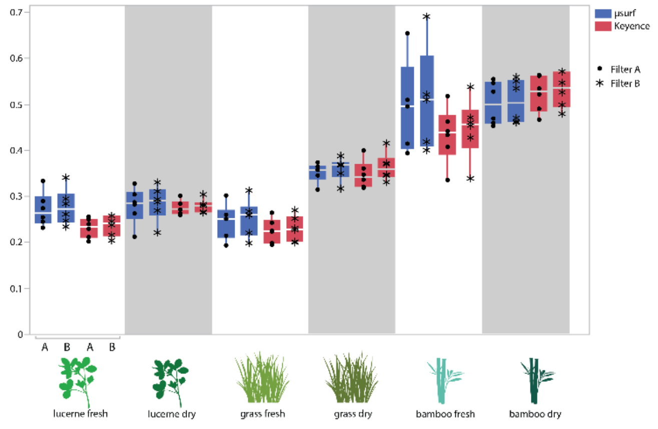


*Vvv*

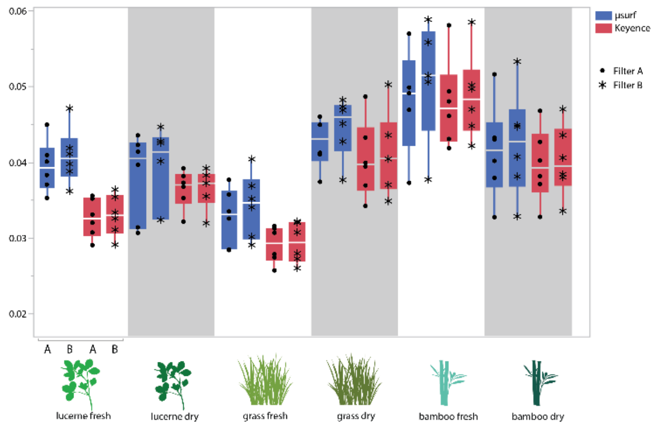


**Figure S1. Boxplots for all 41 analysed dental microwear texture parameters for the upper P4.** Scans captured on Keyence VK-9700 are colour coded in red, scans from $\mu$surf Custom in blue. Filter routine A (gauss filter) is marked with filled circles, filter routine B (median filter) with asterisks. The horizontal bar represents the median; the box encloses the first (25%) and third (75%) quartiles; the whiskers extend to the full interquartile range. For parameter descriptions, see Table S1.

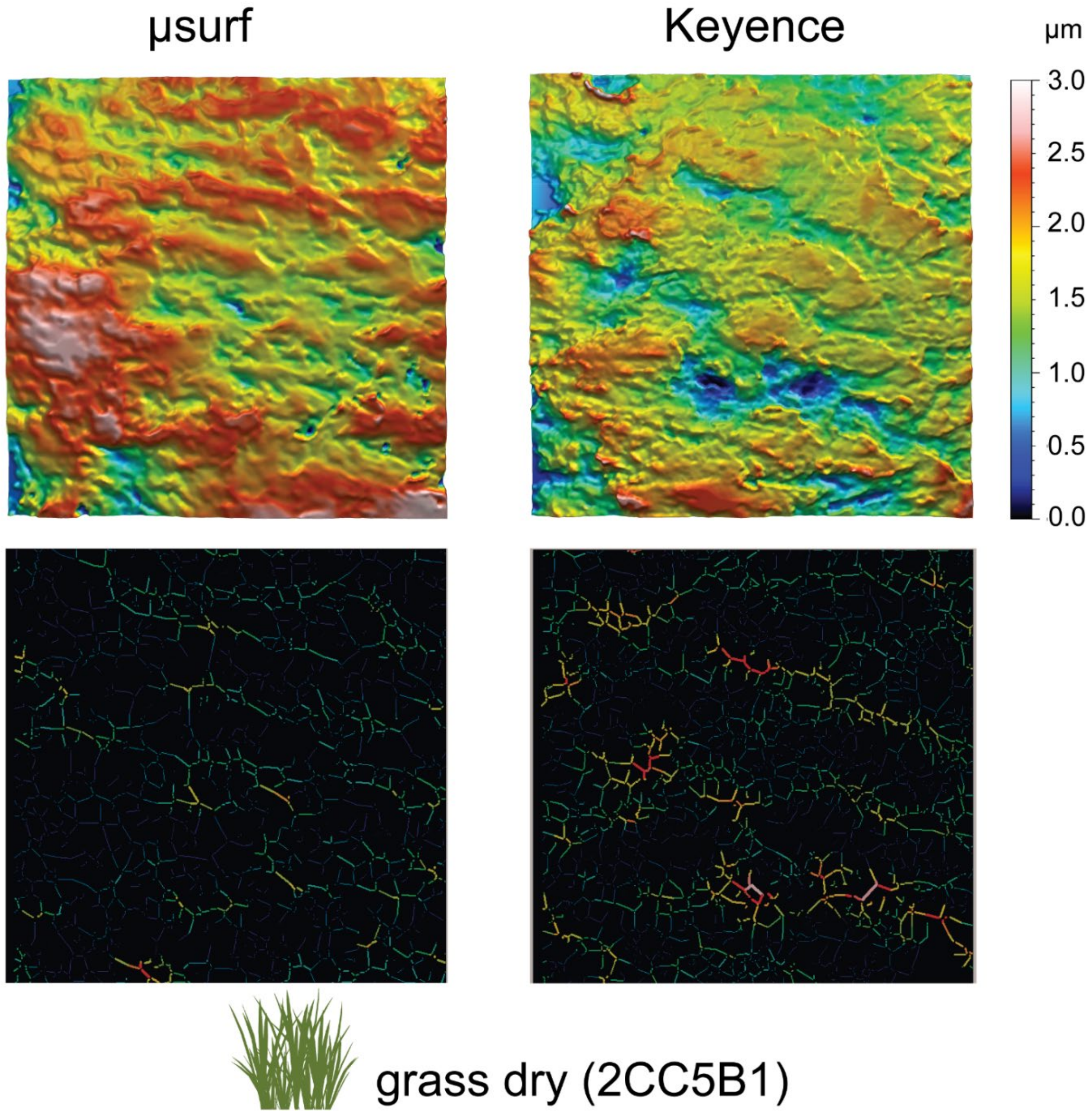


**Figure S2. Exemplary comparison of scans obtained from *µ*surf Custom (left) and Keyence VK-9700 (right).** Upper part shows 3D photosimulations to the same scale. Note the finer grained, sharper structures and peaks on the scan obtained from Keyence VK-9700. Lower part shows furrow maps for the corresponding scans. For the scan from *µ*surf Custom, less furrows are detected. The applied filter routines are combination Keyence* Filter A and *µ*surf* Filter B.
